# Kinematics-based assessment of reaching and grasping movements in LRN ablated animals identifies a role for the LRN in endpoint stabilization and reach timing

**DOI:** 10.64898/2026.04.08.717247

**Authors:** Gavin Thomas Koma, Joshua D. Ross, Thomas J. Campion, Jacquelynn Rajavong, George M. Smith, Andrew J. Spence

**Affiliations:** Temple University, College of Engineering, Philadelphia, USA; Lewis Katz School of Medicine, Medical Education and Research Building, Philadelphia, USA; Department of Pathology and Laboratory Medicine, Children’s Hospital of Philadelphia, Philadelphia, USA

**Author notes:** Asterisks are indicative of co-authorship (Gavin Thomas Koma, Joshua D. Ross).

**Keywords:** Kinematics, Reaching, Lateral Reticular Nucleus

## Abstract

The lateral reticular nucleus (LRN) is thought to contribute to skilled forelimb control but its specific contributions to reaching and grasping remain unclear. In this paper, we examine skilled reaching in intact adult female Long-Evans rats after bilateral LRN ablation through single-pellet reaching tasks. Tasks were analyzed using sensitive quantitative kinematic analyses and qualitative behavioral scoring. Overall, limb transport was largely preserved after ablation, with results appearing in temporally restricted differences. The clearest deficits emerged in pellet-directed endpoint control. LRN-ablated animals showed broad variability in end-point covariance, endpoint spread, and increased trial-to-trial variability, indicating that the movement became less precise and less consistent. These effects were more consistent than any single spatial difference seen, suggesting that ablation of the LRN impairs movement refinement rather than inducing a simple directional bias, although the paw height during the reach was significantly effected. Reach duration also changed, but this temporal difference emerged later and was less prominent. Our results suggest that the LRN acts as an important contributor to endpoint stabilization and reach timing during skilled forelimb behavior.

## Introduction

Dexterous forelimb movements such as reaching, grasping, and food retrieval depend on the coordinated control of the proximal limb during fine digit positioning Whishaw et al. (1992). These behaviors are acquired and refined through practice and are essential for functional interaction with the environment for many species. Despite their importance, the neural circuit mechanisms that support the execution and refinement of skilled forelimb movements remain incompletely understood Krakauer et al. (2019); Klein et al. (2012); Bova et al. (2021). An improved understanding of these mechanisms is important not only for defining how motor skills are learned under normal conditions, but also for identifying internal circuits that may be targeted to improve motor recovery after neurological injury Krakauer et al. (2019).

Rodent skilled reaching is a contemporary model used for studying dexterous motor control Klein et al. (2012). Reach-to-grasp movements in rats share a variety of key organizational features with those observed in primates and humans, which makes this task ideal for investigating the neural basis of skilled motor behavior Whishaw et al. (1992); Klein et al. (2012). In addition, rodents are generally not innately proficient at single-pellet retrieval, but instead improve gradually over repeated behavioral training sessions. This makes the task well suited for studies focused not just on motor performance but also motor learning Bova et al. (2021). Historically, behavioral assessments in a skilled reaching paradigm relied heavily on reach success measures (i.e., successfully obtaining the pellet of food) Klein et al. (2012); Bova et al. (2021); Parmiani et al. (2019). While useful, such measures only provide limited information regarding the organization, refinement, or disruption of the movement itself Klein et al. (2012); Parmiani et al. (2019). Several scoring systems have been developed to evaluate individual movement components, but these approaches remain labor-intensive, subjective, and may not be sensitive enough to fully capture subtle changes in movement structure Whishaw et al. (2008); Mathis et al. (2018); Nath et al. (2019).

Recent advances in pose estimation and kinematic analysis have enabled a more detailed characterization of skilled reaching behavior Mathis et al. (2018); Nath et al. (2019); Bova et al. (2021); Parmiani et al. (2019). Such methods have revealed that gross features of forelimb movement, inclusive of trajectory, end-point variability, and duration, for example, improve within training sessions while other features may evolve over an even longer, multiple day timescale. These findings suggest that distinct circuit mechanisms may contribute differently to skilled forelimb movements and their related precise grasping movements Bova et al. (2021); Nica et al. (2018). A more complete understanding of skilled forelimb control may therefore derive from detailed kinematic analysis of the features that underlie successful performance Nica et al. (2018); Parmiani et al. (2019); Klein et al. (2012).

One circuit of particular interest in this context is the lateral reticular nucleus (LRN) and its associated propriospinal and cerebellar connections Alstermark and Ekerot (2013, 2015); Jiang et al. (2015). The LRN is a precerebellar nucleus positioned to relay motor-related signals to the cerebellum. Here, they likely are integrated with incoming sensory feedback to support online movement correction Alstermark and Ekerot (2013, 2015); Azim and Alstermark (2015). Within the cervical spinal cord a specialized population of C3-C4 propriospinal neurons project both to forelimb-related motor circuitry and to the LRN, thus, forming an anatomical substrate through which descending motor commands may be conveyed as an internal copy of the motor plan Azim et al. (2014b); Azim and Alstermark (2015). These propriospinal neurons receive convergent input from many varying sources: the corticospinal tract, rubrospinal tract, reticulospinal tract, and many other descending systems. Due to this, it is likely that these propriospinal neurons participate in the control and calibration of skilled forelimb movements Azim et al. (2014b); Azim and Alstermark (2015).

Functional studies support that the LRN pathway likely contributes to forelimb motor functions Azim et al. (2014b); Kinoshita et al. (2012); Alstermark and Ekerot (2015). Manipulation of the C3-C4 propriospinal neurons have been shown to produce errors in forelimb kinematics and disruption of LRN-associated circuitry has been linked to deficits in reaching and grasping behaviors Azim et al. (2014b); Kinoshita et al. (2012); Azim and Alstermark (2015). Together, these observations have led to the proposal that the C3-C4 propriospinal-LRN-cerebellar pathway contributes to an efference copy mechanism that supports real-time error correction during movement Jiang et al. (2015); Azim et al. (2014b); Azim and Alstermark (2015); Alstermark and Ekerot (2015). Therefore, it is likely that motor-related signals that are transmitted via the LRN are compared against ongoing sensory information within cerebellar circuits, allowing for corrective signals to be relayed back to descending motor systems Alstermark and Ekerot (2013, 2015); Azim and Alstermark (2015); Huma and Maxwell (2015). Although this model is anatomically and physiologically compelling, the precise contribution of the LRN to skilled reaching and grasping remains insufficiently tested *in vivo* Alstermark and Ekerot (2013, 2015); Jiang et al. (2015).

In addition to contributing to active error correction, the LRN may help signal progression through the movement, providing information that could regulate the timing or gating of successive reach and grasp components. Prior work has shown that perturbation of propriospinal and LRN-associated pathways produces errors in reaching kinematics and forelimb patterning consistent with impaired movement calibration Azim et al. (2014b); Azim and Alstermark (2015); Azim et al. (2014a); Kinoshita et al. (2012). Disruption within this pathway is expected to interfere with the spatiotemporal organization of reaching and grasping rather than simply eliminate movement outright because the LRN relays integrated motor information to cerebellar circuits involved in movement coordination Alstermark and Ekerot (2013, 2015); Thanawalla et al. (2020); Rand et al. (2000). Even with contemporary physiological evidence linking the LRN to skilled forelimb control, the specific consequences of direct LRN ablation have not been defined. In particular, it is unclear whether the loss of function in the LRN primarily disrupts gross forelimb trajectory control, fine grasp-related movements, or the coordination between these components. Addressing this question requires quantitative behavioral analysis capable of resolving detailed kinematic changes beyond simple measure of pellet retrieval success Klein et al. (2012); Mathis et al. (2018); Nath et al. (2019); Bova et al. (2021); Parmiani et al. (2019).

In the present study, we examined the role of the lateral reticular nucleus in skilled forelimb control in intact adult female Long-Evans rats using a bilateral, cell type-targeted ablation strategy in vGlut1-Cre transgenic animals. To define the behavioral consequences of LRN loss, we paired the single-pellet reaching task with a multi-level analysis of performance that included pellet retrieval success, qualitative reach scoring, and quantitative kinematic measurements of forelimb movement patterning. We specifically assessed features of reach organization that are not captured by endpoint success alone, including end-of-reach covariance structure, forelimb trajectory shape, reach duration, and trial-to-trial variability, thereby enabling a detailed evaluation of how LRN ablation alters skilled movement execution. Given the proposed role of the LRN in relaying motor-related signals to cerebellar circuits involved in movement calibration, we hypothesized that bilateral LRN ablation would produce measurable abnormalities in the pattern, consistency, and precision of skilled reaching movements. The objective of this study was therefore to determine how the LRN functions to aid in normal skilled forelimb kinematics and to define the specific aspects of reaching behavior that depend on its integrity.

## Materials and Methods

### Subjects

The rats used in this study were engineered to have a Cre knock-in at the vGlut1 (SLC17A7) locus. This was after our own RNAscope investigation that demonstrated that the neurons of the LRN express vGlut1 and do not express GABA (suggesting their excitatory nature), and that there are very few if any other parts of the caudal medulla containing neurons which express vGlut1, none of which are close enough to the LRN to be of concern for our particular injection surgery. Two litters of heterozygous vGlut1-Cre Long Evans rats were bred, using two female homozygous vGlut1-Cre rats and two male wild types (initial *n* = 18). Their genotypes were confirmed via gel electrophoresis by gathering tail snips, digesting them, purifying the DNA and running PCR using the primer sets provided with the transgenic rats (Rat Slc17a7(Cre) Knock-In, LFKIR-211020-CHG-01, Cyagen, Santa Clara, CA). These rats were weaned at P21 and placed into paired housing based on sex.

### Pre-Operative Behavioral Assessment and Reaching Training

The baseline motor skill of these rats was measured using IBB, grooming, and ladder walk assays at P54 Al-Nasser (2014); Maze Engineers, Conduct Science (2019); Metz and Whishaw (2009). The rats then began reaching training at P71, which was comprised of three weekly sessions (MWF) of 15-20 attempts. The recordings for kinematic analysis began at P108 and were done on the Friday of each week while training was still being performed Mondays and Wednesdays. These initial recordings can be considered the Pre-Operative Period. Rats at this stage were excluded if they were poor learners of the task (performed below 30% success rate and/or less than 5 attempts per session), leaving us with *n* = 6 which aligned well with previous cohorts.

### Lateral Reticular Nucleus Injection Surgery

This training and recording in the Pre-Operative Period continued until P141, at which point the 6 remaining rats were randomized into two groups: Experimental (*n* = 4) and Control (*n* = 2). The two rats selected as controls would go on to receive a ‘sham’ injection of AAV2-hSyn-mCherry (Deisseroth (2018)) while the experimental rats received AAV2-mCherry-FLEX-DTA (Uchida (2014)). This was done as a control for the physical effects of the surgery itself (e.g., passage of the needle through the cerebellum), and of transduction of AAV2 and expression of mCherry into the area of the medulla which hosts the LRN. Both viruses were prepared in-house within HEK-293T cells from plasmids purchased on Addgene.

The surgery was performed twice, with the first targeting the left LRN and the second targeting the right LRN. These surgeries were separated by a week due to the increased fatality rate when performing injections into both sides of the ventral medulla in a single surgical session (see Figure 1). We believe this is due to the proximity of the LRN to other portions of the reticular formation medullary nuclei which are responsible for autonomic functions including vasomotor tone; thus, it is plausible that even a slight physical displacement of those nuclei on both sides of the medulla can be deadly. This week-long rest between the injections allowed the areas around the injection site to recover, ensuring less risk for the bilateral surgery (our fatality rate decreased from ∼40% (bilateral injections, same day) to 0% (single side injections, separating by a week). Another improvement to our method of injection was a slower injection rate and a decreased volume of injection than our initial pilot attempts at the surgery. The coordinates used for these injections were as follows:

**Figure 1.**
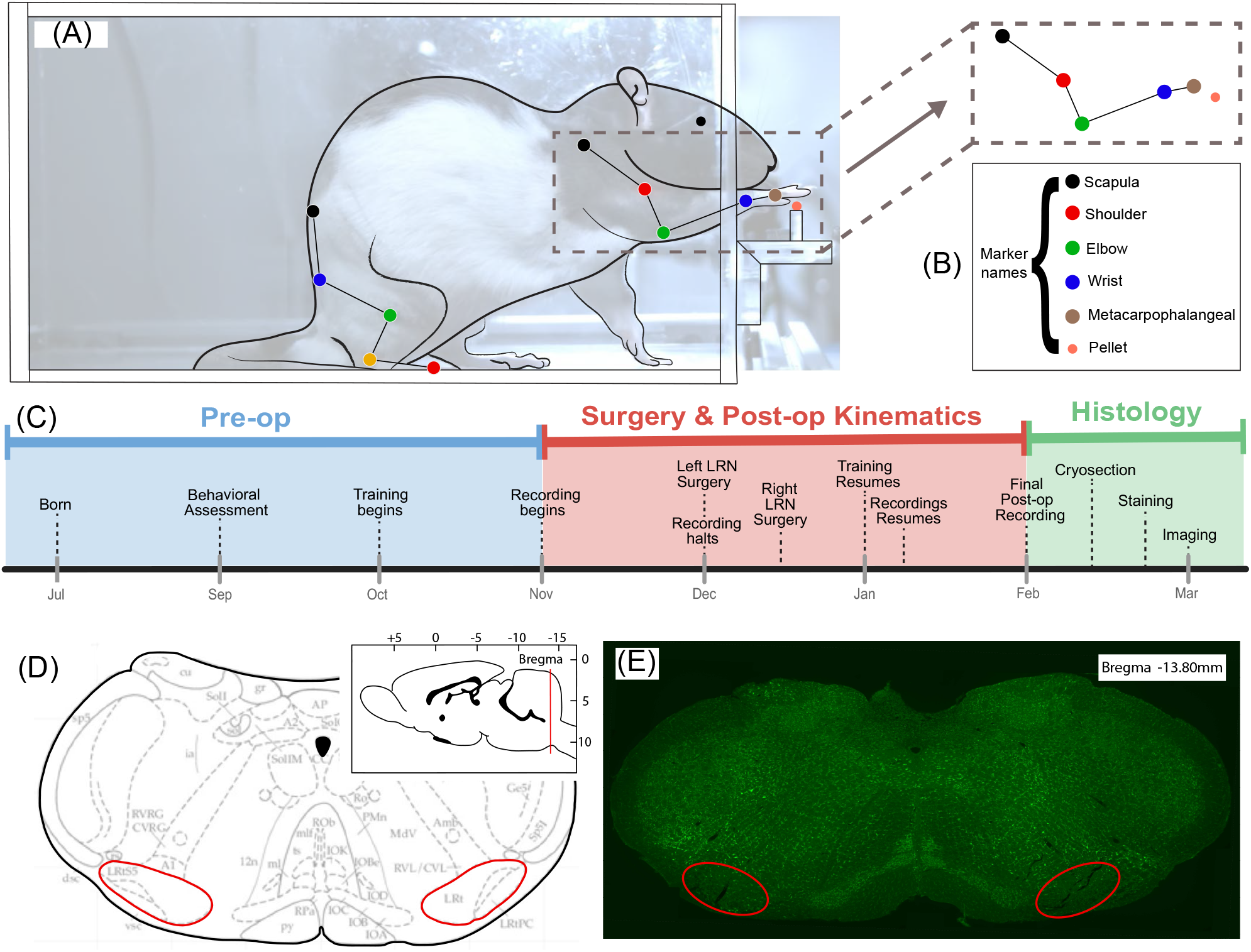
Apparatus, kinematic features tracked, experimental timeline, anatomical location of the LRN and histological confirmation of targeting. (**A**) Representative lateral view of a rat performing the pellet-reaching task, with the tracked forelimb and body landmarks overlaid. (**B**) Marker set used for kinematic reconstruction, including scapula, shoulder, elbow, wrist, metacarpophalangeal joint, and pellet location with hindlimb markers displayed but not analyzed in this study. (**C**) Experimental timeline showing the pre-operative period, bilateral LRN surgeries, post-operative kinematic recording period, and subsequent histological processing. (**D**) Schematic coronal atlas section illustrating the bilateral LRN target region (red outlines contain subnuclei of the LRN: the parvocellular (LRtPC), magnocellular (LRtMC/LRt), and subtrigeminal (LRtS5) divisions), with inset indicating the rostrocaudal level used for histological verification. Atlas adapted from Paxinos and Watson (2013) (**E**) Representative coronal histological section showing bilateral lesion sites within the LRN (red outlines), confirming the location of the targeted ablations. Green stain corresponds to Nissl which stains the endoplasmic reticulum of neuronal cell bodies. Note the lack of green indicates successful ablation of cells in the corresponding areas.

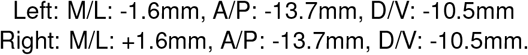

These coordinates are located directly in the center of the LRN respective to all three directional axes. We elected for a single injection in this location due to the high efficiency of the spread of the virus across the whole nucleus without a great deal of leakage to surrounding areas.

The surgical procedure followed a standard stereotaxic brain injection. Rats were first placed under temporary anesthesia with 4% concentrated isoflurane (Isoflurane USP, 78949580, Pivetal, Patterson Veterinary Supply, Loveland, CO) in oxygen. While briefly unconscious, they were weighed and dosed by weight with 5mg/kg Carprofen (Carprofen Injectable Solution, 50mg/mL, 059150, Covetrus, Portland, ME) and a longer term injectable anesthetic mix at 0.1mL/kg (90/10 Ketamine/Xylazine solution; Ketaset, Ketamine HCl Injection, 100mg/mL, KET-00002R3, Zoetis, Parsippany-Troy Hills, NJ; Xylazine, AnaSed LA Xylazine Injection, 100mg/mL, 28745, Vet One, MWI Animal Health, Boise, ID) to last the entirety of the approximately 30-minute-long surgery. Once rats were determined to be fully under anesthesia using paw reflex tests, they were prepared for surgery. The sterile preparation for our subjects included shaving the entire top of the head from the nose all the way back to the base of the skull to expose bregma and to locate the injection site at the caudal most point of the skull. Eye lubrication was placed in the rats’ eyes to prevent formation of cataracts due to dryness out while under anesthesia. In line with proper surgical protocol, three cycles of betadine and 70% ethanol were swabbed liberally across the entire scalp and surrounding area, ensuring that the surgical window was sterile. Rats were then placed carefully onto the stereotaxis by inserting the left ear bar first, then centering the head while placing the right ear bar in. We used a 10*µL* needle (Hamilton #801 Microliter Syringe, 84852, Hamilton Company, Reno, NV) for our injections, pre-filled with the amount of virus required for only the current surgery to avoid thawing the virus prior to use. The needles were aspirated several times with 0.1% sodium dodecyl sulfate (SDS) and 70% ethanol followed by sterile saline and were cleaned between injections as well to ensure sterility. This was continuously lowered until arriving at the dorsoventral location of the LRN, −10.5mm from bregma (lowered slowly over one minute). The needle was lowered an additional 0.1mm and then moved back up to −10.5mm to create a small pocket for actively diffusing virus. A single injection into the nucleus was carried out, with the volume of virus injected into each side 1.5*µL*, over 15 minutes (150*nL* per minute for 10 minutes, with 5 minutes of wait time in between each injection, to allow for dispersion of the virus remaining on and around the tip of the needle). Injection was performed manually using gentle depressions of the plunger. Closure was done using surgical staples (Reflex Clip System, 9mm Clips CAT 201-1000, World Precision Instruments, LLC, Sarasota County, Florida). This was repeated a week later for the contralateral LRN.

### Aftercare

After their scalps were stapled, rats were placed on a heating pad in an recovery chamber, and their eye protection was reapplied. Rats were given sterile saline as a subcutaneous injection. Once the rats awakened, they were placed back into their cages with dietary supplement and 1mg tablet of Rimadyl per rat (Rimadyl 2 mg/tablet, BioServ, Flemington, NJ) was given. Rats were subsequently checked twice daily for three days after the surgery to ensure healthy recovery and were given the same dietary supplement and Rimadyl amounts each time.

### Post-Operative Behavioral Assessment

After a full week of recovery from the second surgery, rats began training on Reaching and Grasping again at P160. Kinematic recordings began on this week, and the schedule returned to training Mondays and Wednesdays and recording Fridays. One rat was dropped from the experimental group due to development of an extremely irregular and untraceable reaching pattern, resulting in the final subject count being three experimental rats and two controls (n=5). The training and recordings were repeated until the rats were P216.

Manual scoring of each training and recording session was performed independently of the kinematics. This was done by marking all reaches; including attempts, successees, and failures of the rats to reach and grasp the pellet, including many which were not recorded and analyzed for kinematic analysis. This yielded comprehensive behavioral scoring data, including the overall successful pellet retrieval rate and overall reaching skill level, scored observationally on a 1-10 scale measure of reaching ability. Their motor skill was once again measured post-operatively using IBB, Grooming and Ladder Walk assays at P222 to confirm whether or not general motor function was affected by our manipulation.

### Automatic 2D Marker Tracking Using DeepLabCut

DeepLabCut (DLC; version 2.1.8.2) was used to estimate the locations of the metacarpophalangeal joint (MCP), wrist, elbow, shoulder, scapula from video recordings. Hindlimb joints were tracked for future analyses. Markers were successfully tracked using similar methodology as described by Mathis et al. (2018) and Nath et al. (2019). Forelimb joints were initially labeled manually and subsequently refined through the iterative correction of model predictions, resulting in a training dataset comprising of 10,550 frames. Video frames had a resolution of 2048×700 pixels. For network training, 95% of the dataset was allocated to the training set and 5% to the validation set using a ResNet-50-based architecture He et al. (2015); Insafutdinov et al. (2016). A p-cutoff of 0.7 was applied to assess the reliability of joint marker detection. The finalized DLC model was then used to automatically track all remaining videos and a custom made application was used to manually clean any jumping or swapping of markers in the final iteration. From these recordings, 163 reaches were extracted and analyzed, corresponding to 163,000 frames. To maintain a conservative statistical framework, data were averaged to a single mean for each rat × treatment ×week except for Figure 5 which compares each individual trial.

### Three-Dimensional Reconstruction and Kinematic Analyses

A custom python code base was utilized to process output from DLC into kinematic variables suitable for statistical analysis Maghsoudi et al. (2017). Two-dimensional joint positions from each camera view were combined with calibration matrices generated through DLTcal in MATLAB (R2025a) to reconstruct three-dimensional joint trajectories. Individual reaching trials were segmented using behaviorally relevant events identified from trial metadata, including paw lift-off, passage through the slit, and trial end. These events were used to define the temporal structure of the reach and to align movements across trials for comparison.

Derived kinematic variables were computed from the 3D trajectories and event-aligned reach segments. These included spatial features of the forelimb trajectory, joint positions across the reach, endpoint distributions, and joint angle measures where relevant. Variability across trials was quantified using standard deviation and covariance measures, allowing assessment of both con-sistency and spread of reach trajectories. For time-resolved analyses, segmented reaches were interpolated to a normalized time base so that kinematic features could be compared across trials and animals despite differences in absolute movement duration.

Standard reaching-related kinematic measures were extracted from these reconstructed trajectories. These included forelimb movement relative to the pellet, vertical and lateral trajectory extent, and endpoint position at key phases of the movement. Joint trajectories and derived measures were then used to quantify changes in reach organization, trial-to-trial variability, and group differences between control and LRN-ablated animals.

### Statistical Analyses

Three-dimensional kinematic coordinates were reconstructed from raw 2D DeepLabCut output using using a custom Python analysis pipeline Maghsoudi et al. (2017), and the MATLAB software package TyDLT Hedrick (2008). Reconstructed 3D kinematic data were processed on a trial-by-trial basis using per-trial summary files together with reconstructed 3D coordinate files. For the principal endpoint and variability analyses, trials were classified via treatment group (control or experimental), behavioral outcome (success or fail), and recording day (12 days pre-surgery, 11 days post-surgery, 32 days post-surgery, and 39 days post-surgery). Pellet-centered coordinates were used for endpoint-based analyses. For the MCP endpoint covariance plots, endpoint coordinates were taken from the peak frame associated with each reach-duration result, and 95% covariance ellipses were computed for visualization of group distributions.

For the derived feature analyses, per-trial kinematic features were extracted from 3D data. Trajectory variability was defined as the mean Euclidean distance of an individual aligned trajectory from the condition-average trajectory after temporal resampling to a fixed number of points. Endpoint spread was defined as the Euclidean distance of each trial endpoint from the centroid of its corresponding day × group× outcome condition. Additional endpoint features included pellet-centered end-point positions in the *x, y*, and *z* dimensions. Overall statistical effects for these features were evaluated using linear mixed-effects models of the form ∼*value* ∼ * *group outcome* + *day* + (1|*rat*), with rat included as a random intercept to account for repeated sampling within animals. Here group refers to the categorical predictor of experimental treatment: control (sham injection of mCherry tracer), and experimental (albation of the LRN with DTA). Outcome refers to the categorical variable for success or failure. For descriptive follow-up comparisons, day-wise pairwise Welch *t* -tests were performed, and only significant comparisons were annotated in the figure panels.

For trajectory-over-time comparisons, wrist trajectories were centered on the pellet and resampled to a normalized time base using the paw-liftoff and paw x-maxima time. Group differences in wrist *x, y*, and *z* position across normalized reach time were assessed separately for each recording day using two-tailed statistical parametric mapping (SPM; Python spm1d package) with *α* = 0.05. The current analysis used the reach-level implementation, in which individual reaches served as observations for the SPM comparison between control and experimental groups. Significant threshold clusters were reported for the relevant axis-by-day comparisons. Reach duration was analyzed separately from the endpoint-feature models. For each trial, reach duration was defined from the beginning of the trial window to the frame at which the selected marker reached its minimum *x*-position within that window. Durations were computed in frames and converted to milliseconds using the frame rate of 250 frames per second. Group differences in reach duration were then evaluated independently for each recording day using two-sided Mann-Whitney *U* tests at the trial level. These per-day duration comparisons were treated as exploratory, and significant comparisons were indicated directly on the box-plot panels.

### Perfusion and Dissection

The precision and extent of ablation in the LRNs of our subjects was quantified using histology as follows. Rats were injected with Fatal Plus pentobarbital solution (Covetrus) at a dosage of 0.2ml/kg [double check concentration] as a primary euthanasia method. Rats were exsanguinated and then perfused with paraformaldehyde. Exsanguination was completed with phosphate-buffered water (PB), and perfusion was completed with a 4% paraformaldehyde (PFA) solution in PB. The brain was cut just caudal to the pyramids to separate brain-stem from spinal cord, and the entire brain was removed and placed into a vial containing 4% PFA solution in PB.

### Histology

Brains were post-fixed overnight in 4% PFA and cryoprotected in 30% sucrose for 48 hours, remaining in sucrose until they sank. They were then cut into 3– 5 mm sections using a rat brain matrix. The caudal-most brainstem section was embedded in matrix, frozen slowly on dry ice, and sectioned coronally at 30 *µ*m on a cryostat maintained below *−*20°C. Sections were collected serially into a 6-well plate so that each well contained representative samples across the full section and were stored in cryoprotectant containing PB, polyvinylpyrrolidone, 30% sucrose, and 30% ethylene glycol. Sections selected for staining were transferred to PBS, while the remaining sections were stored at − 20°C.

For immunostaining, sections were washed 3 times for 10 minutes in 0.04% PBS-Tx on a shaker at 100 rpm at room temperature, briefly rinsed in PBS, and blocked for 1 hour in dilution buffer containing 5% normal donkey serum. Sections were then incubated overnight at 4°C in rabbit anti-dsRed primary antibody (1:1000 in 5% NDS blocking buffer). The following day, sections were washed 5 times for 10 minutes in 0.04% PBS-Tx and incubated for 2 hours at room temperature in donkey anti-rabbit Alexa Fluor 594 secondary antibody (1:1000 in blocking buffer). During the final 20 minutes of secondary incubation, NeuroTrace fluorescent Nissl stain was added at a final concentration of 1:500. Sections were then rinsed 3 times for 10 minutes in PBS and mounted with Fluoromount-G on glass slides with coverslips. Imaging was performed using a Zeiss AxioImager M1 with AxioVision software.

Images were generated by taking 6 subsection images with a 5x objective and piecing them together from both the green and red channels using the software. The extent of ablation was confirmed by comparing the amount of signal from the Nissl stain in the area of the LRN in various slices for each experimental rat to the amount of Nissl signal in the control rats, demonstrating the difference between a living (control) and mostly if not entirely dead (experimental) nucleus. Accuracy of the injection site is demonstrated by the low amount of mCherry expression in the areas surrounding the LRN in the experimental rat samples (**Figure 1E**).

## Results

To determine how bilateral LRN ablation affected skilled reaching, we first examined whether experimental animals retained the overall spatial structure of the reach trajectory across recording days. We then asked whether more subtle abnormalities emerged in the organization and consistency of the movement by quantifying endpoint distributions, covariance, and trial-to-trial variability in pellet-centered coordinates. Finally, we assessed whether these kinematic changes were accompanied by alterations in reach timing. This stepwise approach allowed us to distinguish between gross disruption of reach generation and more selective impairments in movement precision and calibration following LRN ablation.

The initial kinematic analyses indicated that bilateral LRN ablation did not abolish the overall transport trajectory of the limb, but instead produced selective and temporally restricted alterations in wrist motion. As shown in Figure 2, pellet-centered wrist position was examined over normalized reach time in the *x, y*, and *z* dimensions across four recording sessions, with panels A-C corresponding to 12 days pre-surgery, panels D-F to 11 days post-surgery, panels G-I to 32 days post-surgery, and panels J-L to 39 days post-surgery. The surgery was to inject the viral construct that was either the control vehicle or the DTA construct that results in ablation of the LRN over the time course of several days. Across these sessions, the mean control and experimental trajectories generally followed similar time courses, and the overall shape of the wrist trajectory was largely preserved between groups despite the emergence of axis-specific differences during discrete portions of the reach. Reach-level statistical parametric mapping (SPM; *α* = 0.05) further supported this interpretation, identifying significant group differences only in a limited subset of comparisons: a brief cluster in the *x*-dimension on the pre-surgical session in panel A at normalized time 0.0000-0.0206 (*p* = 0.0483), a second cluster in the *x*-dimension at 32 days post-surgery in panel G at normalized time 0.3076-0.3439 (*p* = 0.0435), and a broader cluster in the *z*-dimension at 39 days post-surgery in panel L at normalized time 0.1342-0.2662 (*p* = 0.0028); all remaining axis-wise comparisons were non-significant. This pattern argues against a generalized failure of reach execution following LRN ablation. Rather than disrupting wrist transport through-out the entire movement, the lesion effect appeared to emerge only during restricted temporal intervals and in specific spatial dimensions, consistent with a more selective disturbance of movement control.

**Figure 2.**
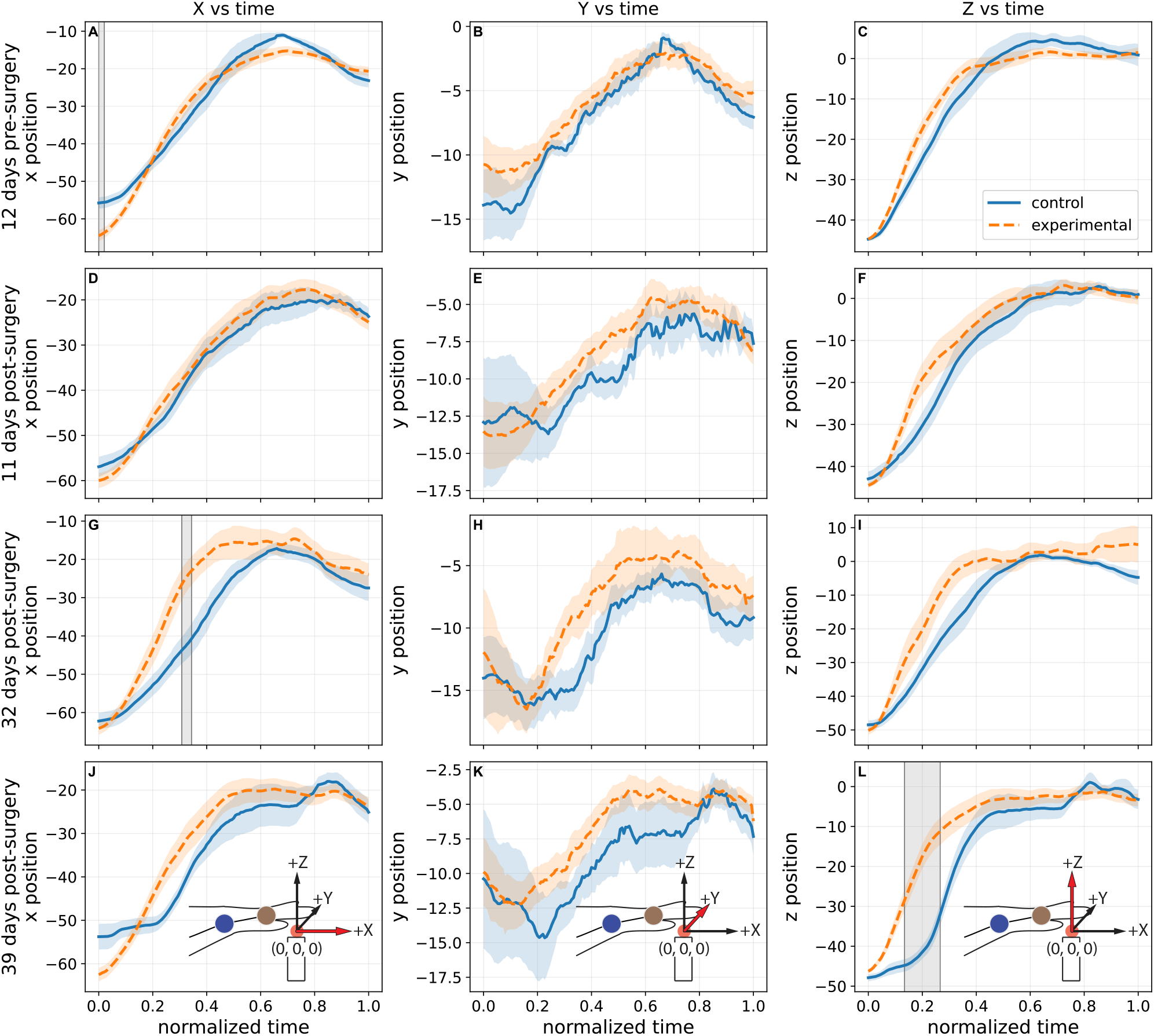
Comparison of Control vs. Experimental wrist 3D trajectories. Panels (**A-C**) correspond to day -12, pre-surgery, (**D-F**) to day +11, (**G-I**) to day +32, and (**J-L**) to day +39, the latter three being post-ablation. Within each date, plots show wrist *x, y*, and *z* position, respectively, against normalized reach time. Please note orientation images in the bottom row illustrating the coordinate system definition relative to the reaching task; highlighted red arrows correspond to the axis of the coordinate being graphed in that column. Curves show the trial-level mean and shaded bonds indicate standard error of the mean (SEM), with each reach analyzed as an individual trial. Across all dates, the overall shape of the wrist trajectory was largely preserved between groups, although axis-specific differences emerged during discrete portions of the reach. Reach-level statistical parametric mapping (SPM; *α* = 0.05) detected significant group differences in *x* on day -12 (**A**; normalized time 0.0000-0.0206, *p* = 0.0483) and day +32 (**G**; 0.3076-0.3439, *p* = 0.0435) and in *z* on day +39 (**L**; 0.1342-0.2662, *p* = 0.0028), whereas the remaining comparisons were not significant. Gray vertical lines denote significant SPM clusters. Rat-averaged results are presented in the supplementary data.

Figure 3 reinforces this conclusion from a spatial perspective. In panels A-D, which correspond to the same dates as in Figure 2, pellet-centered wrist trajectories from control and experimental animals followed broadly similar gross paths toward the pellet origin, with thin lines representing individual trials and thick lines representing group mean trajectories. Although visible day-specific differences in mean trajectory shape and trial-to-trial dispersion were present, the overall 3D organization of the reach remained recognizable in the experimental group across all days. Taken together, Figures 2 and 3 suggest that bilateral LRN ablation left the task of skilled reaching largely intact while introducing modest but statistically detectable deviations in selected dimensions and phases of the movement. This distinction is important for interpretation of the subsequent analyses, because it does indicate that the principle effect of LRN ablation is unlikely to reflect simple loss of reach generation, but instead points towards a more specific impairment in the calibration, consistency, and fine spatial control of the pellet-directed forelimb trajectory.

**Figure 3.**
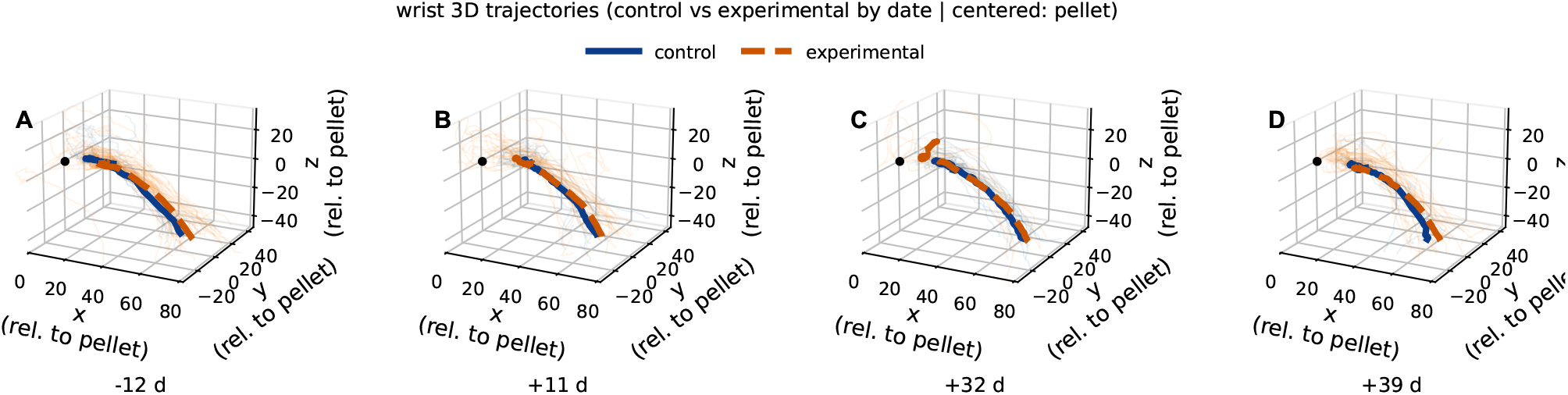
Comparison of Control vs. Experimental wrist 3D trajectories. Panels (**A-D**) correspond to day -12, day +11, day +32, and day +39, respectively. Wrist trajectories are shown in pellet-centered coordinates with thin lines denoting individual trials and thick lines denoting group mean trajectories for control and experimental animals. Across days, both groups followed similar gross reach paths, with visible day-specific differences in mean trajectory shape and trial-to-trial dispersion. This figure provides a spatial visualization of wrist trajectories; formal axis-wise statistical comparisons over normalized reach time are reported in Figure 2. The black point marks the pellet origin. Endpoint varability was notably larger in experimental animals at the +39 time point, seen here as dispersion of individual orange lines.

Subsequent analyses indicated that the more prominent consequences of bilateral LRN ablation emerged not in the gross transport path of the limb, but in the spatial organization and consistency of pellet-directed endpoints. This pattern is illustrated in Figure 4, which shows pellet-centered MCP endpoint distributions across recording days in both the *x*-*y* projection (panels A, C, E, and G) and the corresponding *x*-*z* projection (panels B, D, F, and H). Across sessions, the experimental group frequently exhibited broader 95% covariance ellipses and mean endpoints that appeared more displaced relative to both the pellet origin and the corresponding control distributions. On the pre-surgical session (panels A and B), endpoint spread was already somewhat greater in the experimental group, although both groups remained relatively constrained in space. At 11 days post-surgery (panels C and D), the end-point cloud in the experimental group remained visibly broader than that of controls, suggesting reduced trial-to-trial consistency in where reaches terminated relative to the pellet. This impression persisted at 32 days post-surgery (panels E and F), despite the smaller number of included trials on that day, and was especially pronounced at 39 days post-surgery (panels G and H), where experimental distributions appeared both more scattered and less tightly centered. Because all end-points were plotted in pellet-centered coordinates, this broader and more displaced covariance has a direct functional interpretation: the limb continued to approach the target, but the final placement of the distal forelimb relative to the pellet was less spatially constrained after LRN ablation. Thus, Figure 4 suggests that the principal lesion-related abnormality was not simple failure to launch the reach, but rather diminished precision and reproducibility of the movement as it approached its endpoint.

**Figure 4.**
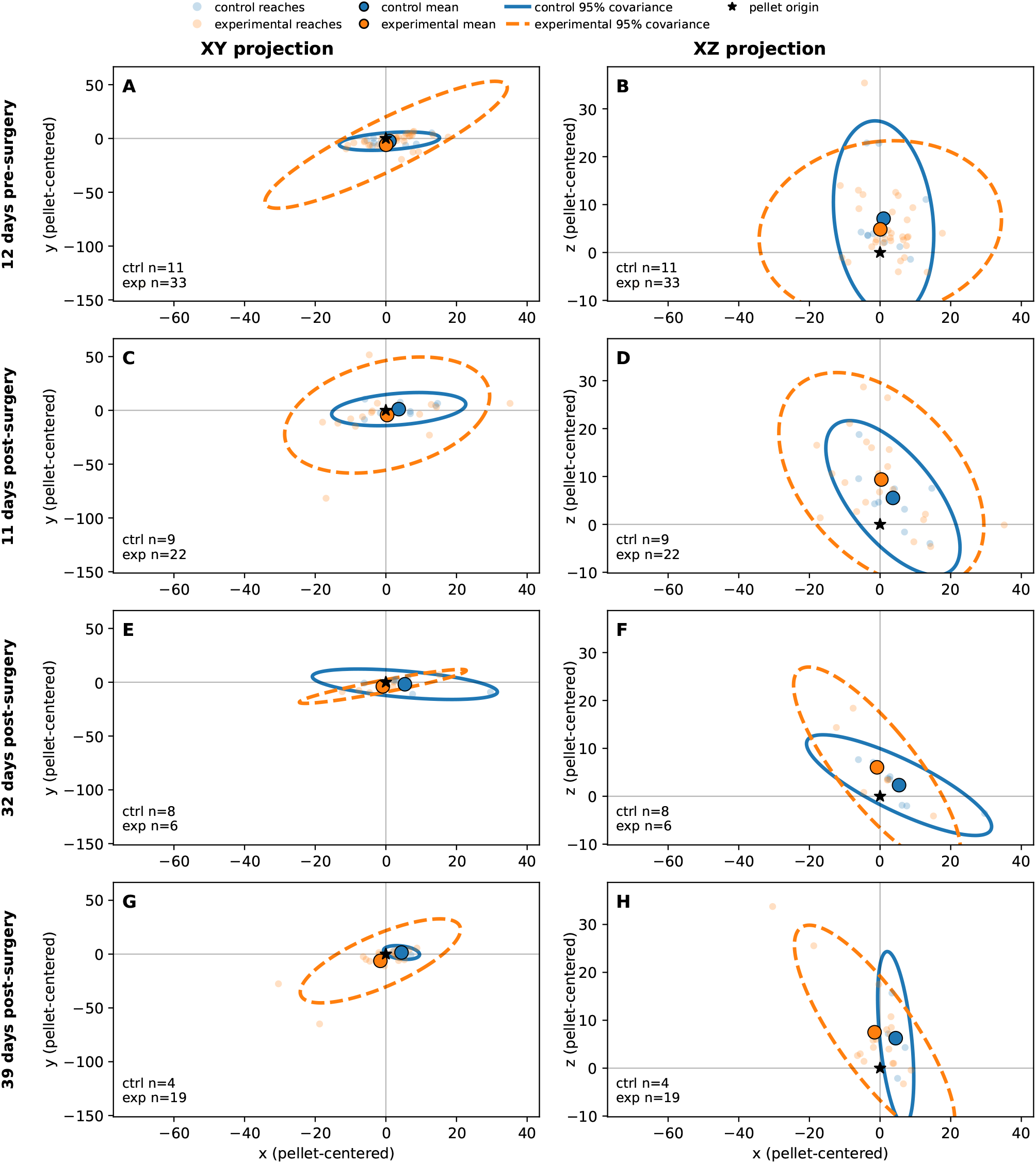
Pellet-centered MCP endpoint covariance in control and experimental animals across recording days. Panels **A, C, E**, and **G** show the pellet-centered *x*-*y* projection of MCP endpoints, whereas panels **B, D, F**, and **H** show the corresponding *x*-*z* projection. The XY projection corresponds to a top-down view of the paw, with +x being a reach towards the pellet through the slit (c.f. axes illustration in **Fig. 2J-L**). The X-Z projection is fore-aft/vertical projection. Rows correspond to four recording days: **A, B**: 12 days pre-surgery; **C, D**: 11 days post-surgery; **E, F**: 32 days post-surgery; and **G, H**: 39 days post-surgery. Small points indicate individual reaches, large filled circles indicate the group mean endpoint for control and experimental animals, and ellipses indicate the 95% covariance region. The black star marks the pellet origin. Across recording days, the position and spread of MCP endpoints differed visually between groups, with experimental animals often showing broader and more displaced endpoint covariance than controls, with a notably larger x variability for experimentals in day +39. Formal statistical comparisons of endpoint covariance are provided in the following figure.

Figure 5 provides quantitative support for the interpretation that bilateral LRN ablation preferentially disrupted movement consistency and endpoint precision rather than the overall execution of the reach. Among the features examined, the clearest overall lesion-related effect was observed for endpoint spread (panel **E**), defined as the Euclidean distance of each trial endpoint from its day ×group× outcome condition centroid.

**Figure 5.**
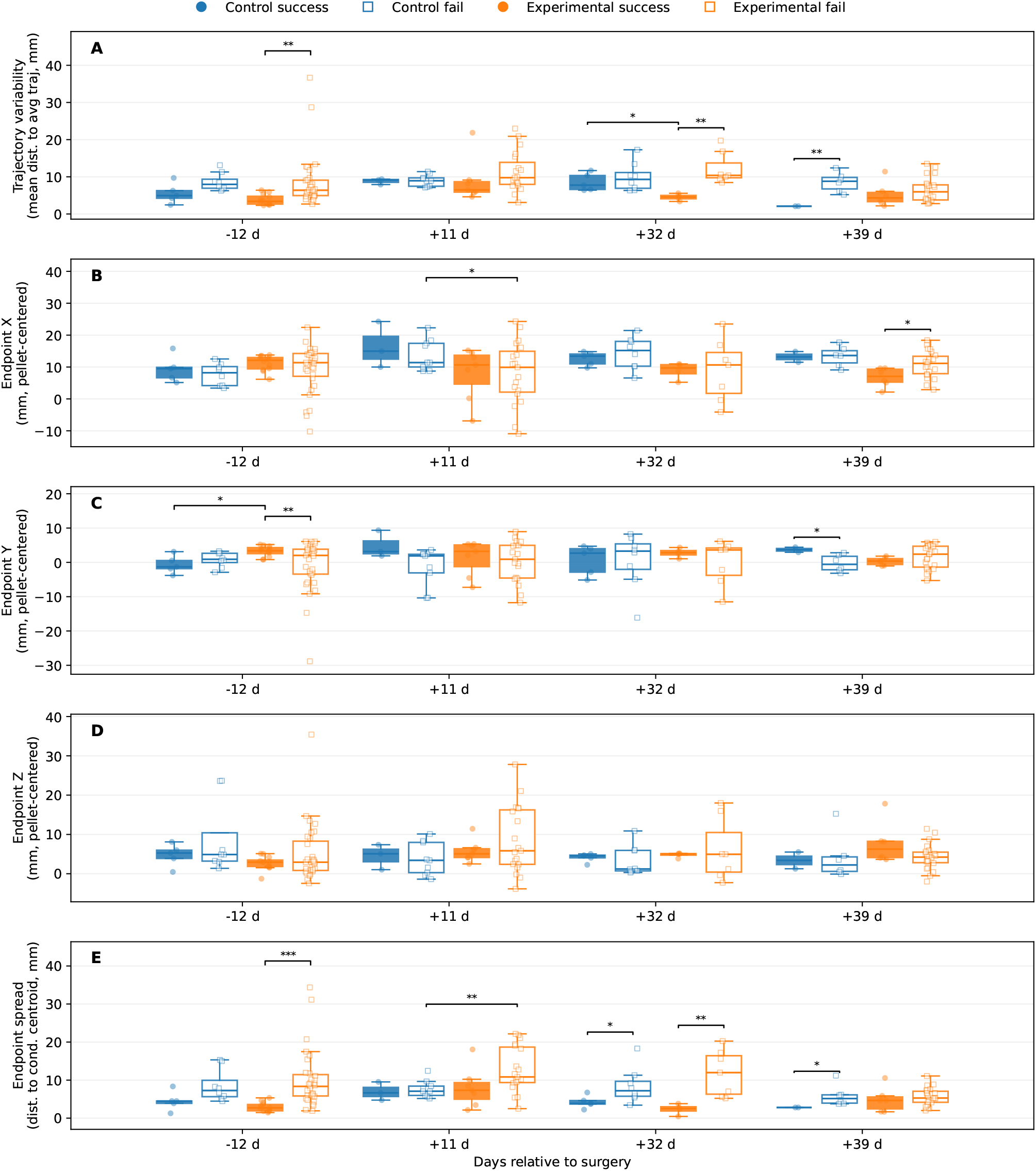
Compiled Feature Analysis of Reaches. (**A**) Trajectory variability, calculated as mean Euclidean distance from the condition-average aligned trajectory. (**B**) Pellet-centered MCP endpoint position in the X dimension. (**C**) Pellet-centered MCP endpoint position in the Y dimension. (**D**) Pellet-centered MCP endpoint position in the Z dimension. (**E**) Endpoint spread, calculated as the Euclidean distance of each trial endpoint from the day × group × outcome condition centroid. For each recording day, box-and-whisker plots with overlaid trial points show control-success, control-fail, experimental-success, and experimental-fail trials. Overall statistics were assessed with mixed-effects models of the form value ∼group *outcome + day + (1| rat) when possible, with OLS fallback otherwise; day-wise brackets indicate significant descriptive Welch post-hoc comparisons only. Trial counts were as follows: on day -12, control fail (*n* = 8), control success (*n* = 5), experimental fail (*n* = 32), and experimental success (*n* = 12); on day +11, control fail (*n* = 9), control success (*n* = 3), experimental fail (*n* = 19), and experimental success (*n* = 7); on day +32, control fail (*n* = 9), control success (*n* = 5), experimental fail (*n* = 7), and experimental success (*n* = 5); and on day +39, control fail (*n* = 6), control success (*n* = 2), experimental fail (*n* = 22), and experimental success (*n* = 6). Endpoint spread (**E**) showed that failures typically ended further from the pellet in general, and at day +11 LRN ablated experimentals had more spread out failures than controls.

Mixed-effects modeling revealed a significant main effect of group for this measure (*β* = 3.2960, *z/t* = 3.1186, *p* = 0.0018), indicating that, across recording days and outcomes, experimental animals exhibited broader end-point dispersion than controls. By contrast, the outcome term (*p* = 0.0662) and the group× outcome interaction (*p* = 0.0705) did not reach significance at the model level, although both trended toward an effect, suggesting that endpoint variability may also have been shaped in part by behavioral success. Day-wise descriptive Welch comparisons were consistent with this interpretation. Within the experimental group, failed reaches showed substantially greater endpoint spread than successful reaches on the pre-surgical day (10.639 1.347 vs. 2.358 ± 0.476mm, *p* = 1.24 ×10^*−*^6), at 11 days post-surgery (13.205 ±1.356 vs. 7.719± 1.947mm, *p* = 0.0389), and at 32 days post-surgery (12.420 ± 2.236 vs. 2.761 ± 0.709mm, *p* = 0.0043). Cross-group comparisons of failed reaches were also significant at 11 days post-surgery (13.205 ±1.356 vs. 6.242 ±1.101mm, *p* = 0.0461), indicating that failed reaches in the experimental group were especially spatially dispersed rela-tive to controls. These finings suggest that the endpoint consequences of LRN ablation were expressed most strongly as reduced clustering of pellet-directed fore-limb placement, particularly under conditions in which reaches were unsuccessful.

A similar, though somewhat weaker, pattern was observed for trajectory variability (panel A), which quantified the mean Euclidean deviation of each aligned reach from its condition-average trajectory. At the model level, neither group (*p* = 0.1200) nor outcome (*p* = 0.0755) nor their interaction (*p* = 0.4925) reached significance, indicating that trajectory-wide variability did not differ uniformly across all conditions. Nevertheless, the day-wise comparisons showed that failed reaches were often less stereotyped than successful reaches, particularly within the experimental group. On the pre-surgical day, experimental failures exhibited greater variability than experimental successes (9.301±1.280 vs. 3.924± 0.420mm, *p* = 0.0003); the same pattern was present at 32 days post-surgery (11.486 ± 1.732 vs. 5.785±0.656mm, *p* = 0.0163). At 11 days post-surgery, failed reaches were also more variable in experimental than in control animals (11.756 ± 1.191 vs. 2.646 ± 0.000mm, *p* = 0.0332). Together, these comparisons suggest that although trajectory variability did not show a uniform global lesion effect, movement was reduced in specific day- and outcome-dependent contexts, again pointing toward instability of reach execution rather than loss of gross movement.

Analyses of pellet-centered endpoint position in the individual spatial dimensions further indicated that lesion-related abnormalities were not distributed uniformly across coordinates. For endpoint *x* (panel B), the overall mixed model revealed no significant main effects or interaction (group: *p* = 0.3351; outcome: *p* = 0.9258; group × outcome: *p* = 0.5487), yet a significant within-experimental outcome difference e was present at 39 days post-surgery, where experimental successful reaches terminated at a more negative *x*-position than experimental failed reaches (-2.097 ±1.334 vs. 1.957 ± 0.902mm, *p* = 0.0302). Endpoint *y* (panel C) showed a comparable absence of effects (group: *p* = 0.4008; outcome: *p* = 0.4060; interaction: *p* = 0.6414), but yielded several notable day-wise differences, including a difference between successful and failed reaches in the experimental group on the pre-surgical day (1.845 ± 0.463 vs. 2.674 ±1.410mm, *p* = 0.0043), a difference between successful and failed reaches in the control group at 11 days post-surgery (4.137±1.137 vs. 0.748±1.370mm, *p* = 0.0269), and a cross-group difference at 39 days post-surgery (*−*0.638 ± 0.626 vs. 1.978 ± 0.266mm, *p* = 0.0085). Endpoint *z* (panel D) similarly showed no significant effects (all *p* ≥0.3941), with the only significant day-wise difference occurring among failed reaches at 11 days post-surgery, where experimental animals terminated farther from the pellet in *z* than controls (7.784± 2.038 vs. 2.127± 1.361mm, *p* = 0.0296). Taken together, these results indicate that bilateral LRN ablation did not impose a simple, fixed spatial offset on reach endpoints. Instead, the strongest and most consistent abnormality was an increase in endpoint dispersion, accompanied by selective day- and outcome-dependent shifts in specific endpoint coordinates. Therefore, these findings do support the interpretation that the LRN contributes less to the overall production of the reach trajectory than to the trial-by-trial stabilization and calibration of pellet-directed fore-limb placement.

The reach-duration analysis indicated that temporal changes after bilateral LRN ablation were present, but were more limited and less consistent that the spatial differences identified in the preceding endpoint analyses (see Figures 4 & 5). As shown in Figure 6, reach duration distributions overlapped substantially between control and experimental animals on the pre-surgical session and at 11 days post-surgery, with no significant per-day treatment differences detected at either time point (12 days pre-surgery: *U* = 364.5, *p* = 0.138; 11 days post-surgery: *U* = 177.0, *p* = 0.520). Descriptive statistics were consistent with this. On the pre-surgical day, median reach duration was 384ms in controls and 326ms in experimental animals, while at 11 days post-surgery the medians were 426ms and 438ms, respectively. By contrast, significant group differences emerged at the later post-surgical sessions. At 32 days post-surgery, experimental animals exhibited shorter reach durations than controls (*U* = 128.0, *p* = 0.0253), with median values of 266ms in the experimental group versus 476ms in controls. A similar pattern was present at 39 days post-surgery (*U* = 169.0, *p* = 0.0315), where median reach duration was 298ms in experimental animals compared with 448ms in controls.

**Figure 6.**
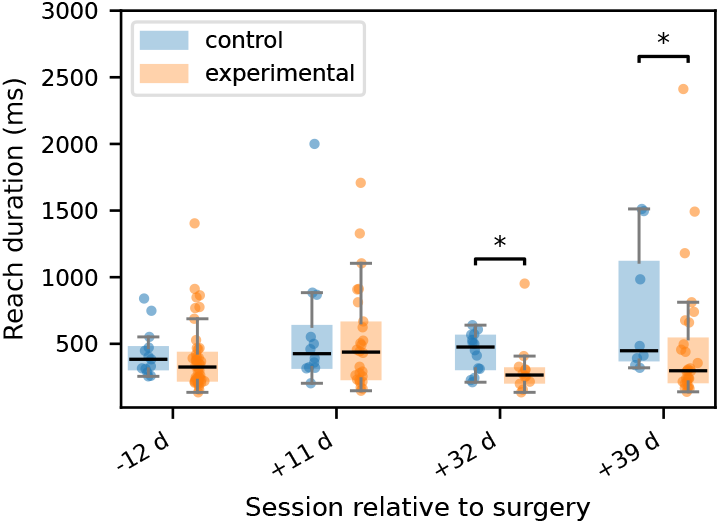
Reach duration across recording day and treatment group. Box-plots show trial-level reach duration for control and experimental animals across recording days, with individual trials overlaid. Reach duration distributions were similar between groups on day -12 and day +11, whereas exploratory day-wise comparisons indicated shorter reach durations in experimental animals on day +32 and day +39. Asterisks indicate two-sided Mann-Whitney U tests performed separately for each day at the trial level; significant differences were observed on +32 (*U* = 128.0, *p* = 0.025) and +39 (*U* = 169.0, *p* = 0.031).

Thus, unlike the earlier trajectory analyses in Figures 2 & 3, which showed broadly preserved reach trajectory and structure with only temporally focused differences, Figure 6 suggests that timing changes were not an immediate result of LRN ablation, but instead emerged later in the post-surgical period.

## Discussion

The present findings indicate that bilateral LRN ablation altered skilled forelimb behavior primarily by degrading the precision and stability of pellet-directed reaching rather than by abolishing the basic transport pattern of the movement. In the trajectory-based analyses, experimental animals retained broadly recognizable wrist paths toward the pellet in both the axis-by-time plots shown in Figure 2 and the 3D trajectory reconstructions shown in Figure 3, with statistically significant group differences limited to a small number of temporally restricted SPM clusters in wrist *x* on day −12 (Figure 2A) and day +32 (Figure 2G), and in wrist *z* on day +39 (Figure 2L); all remaining axis-wise comparisons were non-significant. This relative preservation of kinematic trajectory contrasted with the endpoint-level variance observed in Figures 4 & 5, where experimental animals showed a wider range of MCP endpoint covariance, increased endpoint spread, and greater context-dependent variability, indicating that the final placement of the forelimb relative to the pellet was less spatially constrained after LRN loss. In particular, Figure 4 showed broader and more displaced endpoint covariance in the experimental group across recording days, while Figure 5 identified endpoint spread as the clearest overall lesion-related effect, with a significant main effect of group in the mixed-effects model (*p* = 0.0286). Reach duration, shown in Figure 6, was also altered, but only at the later post-surgical sessions, where experimental animals exhibited shorter reaches than controls at day +32 and day +39, suggesting that temporal differences emerged after, and were less dominant than, the spatial abnormalities. Taken together, these findings support the central interpretation that the LRN contributes less to the generation of the reach in an all-or-none sense than to the trial-by-trial assessment and stabilization of pellet-directed forelimb movement.

The difference between overall trajectory and variability between endpoint organization is informative for interpreting the functional contribution of the LRN. As shown in Figures 2 & 3, experimental animals continued to generate a broadly recognizable pellet-directed wrist trajectory, and formal axis-wise comparisons revealed only a small number of temporally restricted SPM differences rather than a total disruption of the entire reach. By contrast, Figures 4 & 5 showed that the more robust lesion-related phenotype emerged at the endpoint level, where experimental animals displayed broader MCP endpoint covariance and significantly greater endpoint spread than controls. In the mixed effects analysis of endpoint spread, experimental vs. control (group) was the clearest overall predictor, with experimental reaches exhibiting greater dispersion across recording days and outcomes (*p* = 0.0018), whereas the effects of outcome and the group-by-outcome interaction were weaker and did not reach conventional significance thresholds in the MCP analysis. This pattern argues against the interpretation that ablation of the LRN simply caused a fixed spatial offset or a nonspecific loss of movement (reach generation). Instead, it suggests that the basic reaching trajectory remained in tact, while the trial by trial regulation of forelimb placement relative to the pellet became less stable. Functionally, this difference is more consistent with the role of the LRN being primarily involved in refinement, repetition, or adjustment of skilled reaching than in the initiation of the reaching task itself. Thus, the present data suggests that even when the limb could still move toward the target along a general path, the absence of normal LRN contribution was associated with reduced precision in how that movement ultimately brought the forelimb to the pellet.

An additional point emerging from the endpoint analyses is that the effect of bilateral LRN ablation was not expressed exclusively as a shift in group means, but was shaped by trial outcome and by specific kinematic features. In Figure 5, failed reaches were often associated with broader distributions and larger values for both the trajectory variability and the endpoint spread, most particularly within the experimental group, and thereby indicating that the lesion-related disruption became most apparent under conditions in which the behavior was already less successful. For endpoint spread, for example, failed experimental reaches were substantially more dispersed than successful experimental reaches on the pre-surgical session, at 11 days post surgery, and again at 32 days post-surgery. These experimental reaches also exceeded failed control reaches at 11 and 32 days post-surgery. A similar pattern was present for trajectory variability, where failed reaches in the experimental group frequently showed less stereotyped behavior in their trajectories than successful reaches. This information suggests that LRN ablation did not simply impose a single fixed deficit on all reaches. Instead, some reaches remained relatively well organized, whereas others deviated more markedly in end-point precision or trajectory consistency, furthering the hypothesis that LRN ablation does not simply impose a single fixed deficit on all reaches.

This interpretation is strengthened by the fact that the strongest and most consist difference across endpoint analyses was within increased dispersion. In Figure 5, the mixed-effects model for endpoint *x, y*, and *z* did not reveal the same group effect seen for endpoint spread, and the significant day-wise comparisons in individual coordinates were selective. This pattern argues against a simple directional bias in which the LRN ablation caused the forelimb to terminate consistently too far in one direction. Instead, the data are more consistent with reduced repetitive trajectory (i.e., the forelimb trajectory arrived at a broader range of positions relative to the pellet from trial to trial). This outcome-dependent increase in variability suggests that the LRN normally contributes to the reliability with which skilled forelimb trajectories are refined into stable, reproducible pellet-directed endpoints.

Although the most prominent abnormalities after bilateral LRN ablation were spatial rather than temporal, the reach-duration analysis suggests that the temporal organization of pellet-directed behavior was also altered at later post-surgical stages. Reach duration distributions overlapped substantially between control and experimental animals on the pre-surgical session and at 11 days post-surgery, whereas differences emerged at the later post-surgical sessions, where experimental animals exhibited shorter reaches than controls. This temporal pattern appears secondary to the more consistent abnormalities observed in endpoint covariance, endpoint spread, and movement trajectory. Thus, the shorter reaches observed at later post-surgical stages are unlikely to reflect improved motor efficiency, but rather, they may indicate an altered execution strategy in which pellet-directed reaches were completed more rapidly but with reduced consistency. This would corroborate our findings regarding increased endpoint dispersion and a greater trial-to-trial variability observed in the experimental group.

In conclusion, bilateral LRN ablation did not eliminate the ability of a rodent to generate pellet-directed reaches, but instead altered the quality with which those movements were executed. The most consistent abnormalities were observed not in the gross transport trajectory of the limb, but in the precision, reproducibility, and spatial pattern of pellet-directed endpoint control, with additional later-emerging changes in reach timing. This pattern supports the interpretation that the LRN contributes importantly to the calibration and stabilization of skilled forelimb behavior, helping to ensure that the reaches are executed in a consistent manner. More broadly, these findings support the view that disruption of LRN-associated circuitry can degrade dexterous movement by impairing movement refinement and trial-by-trial control, even when the basic structure of the reach remains intact.

## ACKNOWLEDGEMENTS

This work was supported by NIH NINDS grant 1R01NS114007-01A1 to A.J.S., NIH NINDS grant R01NS117749-01 to G.M.S., Shriners Hospitals for Children Grant #85115 to A.J.S., Craig H. Neilsen Foundation Senior Research Grant (#546798) to A.J.S., and the Shriners Viral Core Grant #84051-PHI-21 to G.M.S. We thank Anastasia Baranova for assisting with Panel A figure composition. For the purpose of open access, the authors have applied a Creative Commons Attribution (CC BY) license to any Author Accepted Manuscript version arising.

## AUTHOR CONTRIBUTIONS

GTK, scientific direction, kinematic acquisition, kinematic rig construction, data analysis, programming, wrote the manuscript; JDR, scientific direction, viral preparation, breeding, surgeries, post-surgical monitoring, behavioral training, kinematic acquisition, dissection, histology, microscopy; JR, image/video labeling; AJS, scientific direction, data analysis, programming, editing; GMS, scientific direction, funding acquisition, first draft, editing.

